# Visual Context Influences Lateral Balance While Walking on Winding Paths

**DOI:** 10.64898/2026.01.18.700184

**Authors:** Anna C. Render, Tarkeshwar Singh, Joseph P. Cusumano, Jonathan B. Dingwell

## Abstract

Effectively navigating daily environments necessitates achieving adaptability while maintaining stability. This requires integrating sensory feedback, primarily from the visual system, to maintain balance and maneuverability as we walk. This study examined the impact of visual information salience on lateral balance and stepping. Twenty-eight healthy adults (16F/12M; 26.16±4.23 years) participated. Participants walked along each of two virtual paths (Straight vs. Winding), having each of two path color contrasts (High vs. Low), in each of two environments with differing visual richness (Rich vs. Sparse). We quantified stepping errors as the percentage of steps landing outside designated path boundaries. We computed means (*μ*) and standard deviations (*σ*) of the minimum mediolateral margins of stability (*MoS*_*L*_), and we computed lateral Probability of Instability (*PoI*_*L*_) to assess participants’ risk of taking unstable (*MoS*_*L*_ < 0) steps. On Straight paths, participants made more stepping errors on Low (vs. High) contrast paths for both environments, and exhibited decreased *σ*(*MoS*_*L*_) in Sparse (vs. Rich) environments on paths of both visual contrasts. On Winding paths, participants made the most stepping errors on Low Contrast paths in Sparse environments. They walked with smaller *μ*(*MoS*_*L*_) and exhibited higher *PoI*_*L*_ on Low Contrast paths in both environments, and they exhibited higher *PoI*_*L*_ in Sparse environments on paths of both visual contrasts. The effects of diminished visual information were far more pronounced on Winding paths (vs. Straight), hindering performance and balance maintenance, as these conditions challenged both mechanical and sensory mechanisms underlying balance control.

## INTRODUCTION

Safely navigating our daily environments requires both stability and adaptability [1, 2]. This is especially critical as we often encounter crowded spaces [3, 4], or cluttered environments [5] that challenge our ability to maintain lateral balance as we navigate. Walking involves continuous, dynamic engagement with the environment [6]. Successful locomotion depends on precise coordination between sensory inputs and motor actions, where vision is crucial for identifying hazards, guiding foot placement, and adapting gait patterns to environmental demands [7].

Humans inherently exhibit greater instability in the frontal plane [8]. Conversely, while adaptability allows for precise adjustments to navigate complex environments, it also increases lateral instability while walking [2]. However, adaptability assumes stability as a precondition. To prevent or correct mediolateral instability, individuals primarily rely on adjusting foot placement [9], which serves to redirect their center-of-mass. Despite the vital importance of these adaptive strategies, lateral instabilities and falls remain significant concerns [10], particularly for older adults with impaired vision [11].

People adapt their steps to the walking tasks they perform [2, 7, 12–16]. This in turn, influences their lateral stability. Narrower walking paths lead to more stepping errors (i.e., off-path steps) [14, 17], but only modest increases in the risk of taking unstable steps [14, 18, 19]. Conversely, walking on winding paths can greatly increase both the number of stepping errors and the risk of instability [19].

Walking is a complex balance control process that requires integrating visual, vestibular, and somatosensory feedback, with vision being primary among these [7, 20]. Vision helps guide goal-directed locomotion, thus aiding in stability and orientation, particularly in dynamic environments [7]. Vision allows us to make anticipatory adjustments to gait necessary for safe travel [21]. Visual impairments are well-documented to negatively affect gait parameters and postural control [22], increasing the risk of instability [21].

While previous studies examined how navigating continually-winding paths affects lateral balance [19], the influence of visual characteristics, like the perceptual salience (the quality or prominence of visible features in the environment) [23] of the walking path or surrounding environment, remains poorly understood. Precision stepping tasks have investigated visual influences on stepping accuracy [24], but they are often limited to controlled laboratory environments that do not capture the broader dynamics of real-world path navigation. Thus, it remains unclear how perceptual salience affects lateral instability during path navigation.

Here, we quantified how changes in visual salience affect task performance and lateral balance during walking. Specifically, we manipulated salience by altering the richness of the surrounding environment (rich vs. sparse) and the color contrast of the walking path (high vs. low). Importantly, we kept the biomechanical tasks of walking on straight or winding paths unchanged. This design allowed us to isolate the effects of visual salience on stepping performance and balance control. Extending our previous study that manipulated only path shapes [15, 19], this work integrates sensory aspects of visual guidance to elucidate the interplay of perceptual and mechanical demands on balance control.

We tested two competing hypotheses regarding lateral stability during walking. First, the biomechanical constraint hypothesis posits that stability is primarily governed by biomechanical task demands [6, 25], such that healthy adults would maintain consistent balance regardless of changes in visual salience. Alternatively, the visual dominance hypothesis posits that reduced visual salience might impair balance control independent of the task’s biomechanical demands. This could occur through either or both of two mechanisms. First, if people rely on vision primarily to enhance *task*-relevant information [13, 26] and reduce visual uncertainty [27], then we would expect decreasing visual contrast of the walking paths to degrade balance performance (i.e., increase stepping errors and probability of taking unstable steps). Second, if *peripheral* vision is critical to governing locomotor behavior [28, 29], then we would expect decreasing the visual richness of the surrounding environment to degrade balance performance. By systematically and independently manipulating both aspects of perceptual salience while holding biomechanical task demands constant, this study directly assessed how salience of visual information contributes to both stepping accuracy and balance control during a naturalistic walking task.

## METHODS

### Participants

Twenty-eight adults (16F/12M; age 26.16±4.23 yrs; height 1.73±0.10 m; body mass 70.3±14.9 kg) provided written informed consent, as approved by the Institutional Review Board of Penn State University. Participants were screened to ensure no medications, lower limb injuries, surgeries, musculoskeletal, cardiovascular, neurological, or visual conditions affected their gait.

### Protocol

Participants walked on a 1.2 m wide treadmill in an M-Gait virtual reality system (Motek Medical, Netherlands; Fig.1A). Each participant wore a safety harness. All walking trials were performed at 1.2 m/s. Participants first walked 3-minutes to acclimate to the system.

**Figure 1.**
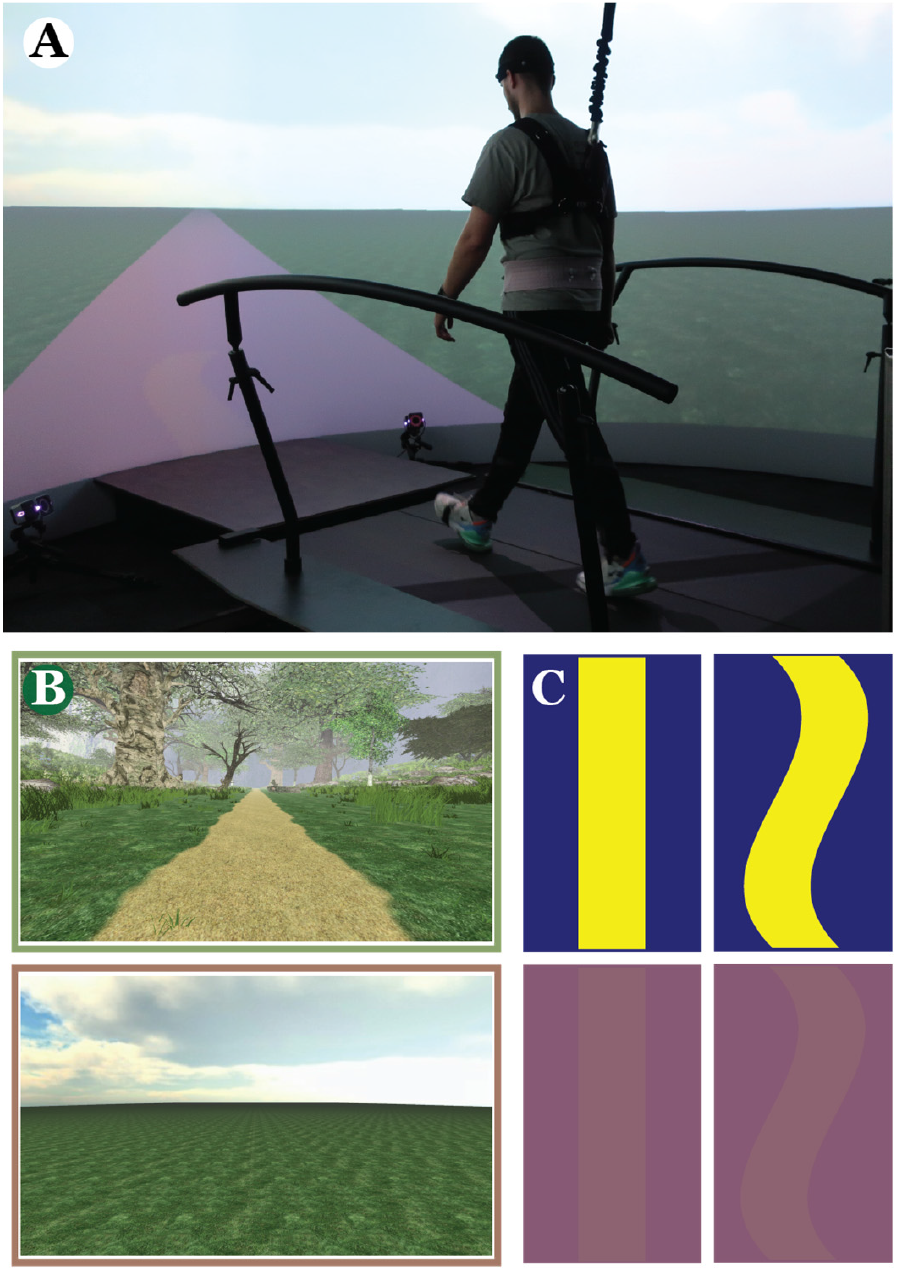
**A)** Photo of a participant walking in the Motek M-Gait system. **B)** Participants walked through each of two virtual environments: a Rich (top) forest scene with dense foliage or a Sparse (bottom) open grassy plain. **C)** In each environment, participants walked on each of four paths, consisting of either high (HC; top) or low (LC; bottom) color contrast paths that were either Straight (STR; left) or Winding (HIF; right). We created the winding (HIF) paths using a sum-of-sines function [15, 19], *z*(*x*) = 0.22sin(1.875*x*) + 0.05sin(2.5*x*) + 0.03sin(3.75*x*), where *z* is lateral position of the path center and *x* is forward distance (both in meters). All paths (STR and HIF) were 0.45 m wide. High Contrast conditions projected a yellow ([R,G,B] = [254,255,0]) path onto a dark blue ([R,G,B] = [0,0,129]) background (top). Low Contrast conditions projected a purple ([R,G,B] = [141,98,113]) path onto a slightly darker purple ([R,G,B] = [134,88,115]) background (bottom).

Participants walked through each of two environments: Rich (Fig. 1B; top) and Sparse (Fig. 1B; bottom). They walked on paths with each of two color contrasts: High (Fig. 1C; top) and Low (Fig. 1C; bottom). Lastly, participants walked on each of two paths: Straight (STR; Fig. 1C; left) and Winding (HIF; Fig. 1C; right). We instructed participants to “stay on the path” and minimize stepping errors. They received visual (firework) and auditory (flute) penalties when steps landed outside the path boundaries.

For each condition (8 total), participants completed a 1-minute introductory trial followed by two 3-minute experimental trials. Order of presentation of conditions was randomly assigned to each participant and counterbalanced across participants using a Latin Square design. Participants were allowed to rest as needed after each trial.

### Data Processing

Each participant wore 17 retroreflective markers: five around the head, four around the pelvis (left and right PSIS and ASIS), and four on each shoe (first and fifth metatarsal heads, lateral malleolus, and calcaneus). Kinematic data were collected at 100 Hz from a 10-camera Vicon system (Oxford Metrics, Oxford, UK) and post-processed using Vicon Nexus software. Marker trajectories and path data from D-Flow software (Motek Medical, Netherlands) were analyzed in MATLAB (MathWorks, Natick, MA).

For each trial, we low-pass filtered (10 Hz 4^th^-order Butterworth) the raw marker trajectories and interpolated these data to 600 Hz. Heel strikes and toe-offs were identified using a previously validated algorithm [30]. For consistency, we analyzed the first *N* = 250 steps of each trial. On the winding paths, one’s direction of motion changes at each new step. We therefore transformed our data into path-based local coordinates [31], where for any given step, the locally lateral *z*-axis was defined to be perpendicular to the path at the midpoint between the left and right foot placements for that step.

### Stepping Errors

We quantified Stepping Errors as the percentage (out of *N* = 250) of steps within each trial where any of the first or fifth metatarsal head or calcaneus markers exceeded the path boundary. This measure quantified overall task performance.

### Lateral Margins of Stability

The lateral margin of stability (*MoS*_*L*_) [32, 33] measures the linear distance from a threshold (i.e., *MoS*_*L*_ = 0) beyond which an inverted pendulum becomes unstable [33]. Here, we quantified lateral stability in the locally-lateral *z*-direction at each step (Fig. 2A). We averaged the motion of the 4 pelvis markers to approximate center-of-mass (CoM) state (*z, ż*) [34]. We then calculated the *minimum MoS*_*L*_ within each step *n*:

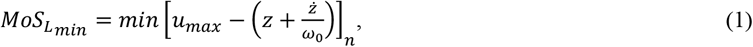

where the *z*-coordinate of the leading foot’s fifth metatarsal marker approximated the lateral boundary of the base of support (*u*_*max*_), and 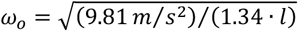, where *l* was leg length [35]. Accordingly, a positive *MoS*_*Lmin*_ is typically interpreted to indicate a “stable” step, while a negative *MoS*_*Lmin*_ is interpreted as an “unstable” step [32, 33].

**Figure 2.**
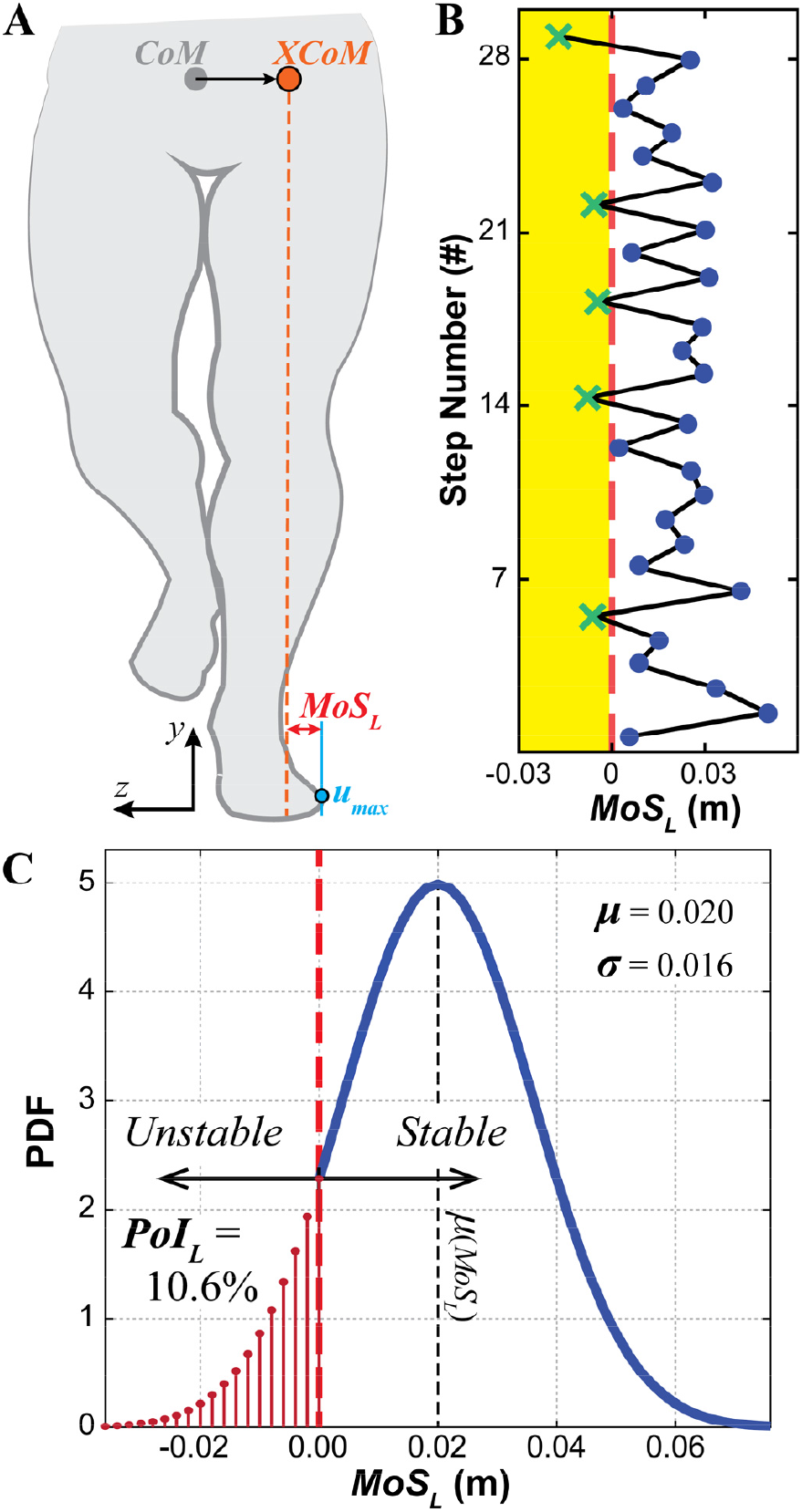
**A**) Schematic showing the variables used to calculate *MoS*_*Lmin*_ (Eq. 1): center-of-mass (CoM; grey), extrapolated center-of-mass (*XCoM*_*L*_ = *z*+*ż*/*ω*_o_; orange), and lateral base-of-support boundary (*u*_*max*_; blue). **B**) Example time series data of *MoS*_*Lmin*_ across several steps. The vertical red dashed line indicates *MoS*_*Lmin*_ = 0. Stable (*MoS*_*Lmin*_ > 0) are shown as blue dots (●). Unstable (*MoS*_*Lmin*_ < 0) steps are shown as green crosses (×). **C**) Hypothetical probability density function (PDF) to demonstrate how to calculate *PoI*_*L*_. The dashed vertical red line at *MoS*_*Lmin*_ = 0 separates stable (*MoS*_*Lmin*_ > 0) from unstable (*MoS*_*Lmin*_ < 0) steps. Note the mean, *μ*(*MoS*_*Lmin*_) = 0.020 m, is well within the *stable* region, and thus unrelated to the *un*stable steps. Conversely, *PoI*_*L*_ (Eq. 2) directly calculates the cumulative probability precisely of those *un*stable (*MoS*_*Lmin*_ < 0) steps. Hence, *PoI*_*L*_ is consistent with Hof’s theory, whereas *μ*(*MoS*_*Ln*_) is not.

For each trial performed by each participant for each condition, we extracted time series of *MoS*_*Lmin*_ (e.g., Fig. 2B) from all *n* ∈ {1, …, 250} steps. We computed within-trial means, *μ*(*MoS*_*Lmin*_), and standard deviations, *σ*(*MoS*_*Lmin*_), for each trial.

### Lateral Probability of Instability

However, averaging *MoS*_*L*_ across steps cannot properly estimate risk of becoming unstable [18]: i.e., the likelihood that any given step will exceed the *MoS*_*L*_ = 0 threshold defined by Hof [33]. We therefore calculated lateral *Probability of Instability* (*PoI*_*L*_) [18], which specifically estimates the cumulative probability (P) that steps within an assumed normal distribution of *MoS*_*Lmin*_ values will exceed *MoS*_*Lmin*_ < (Fig. 2C):

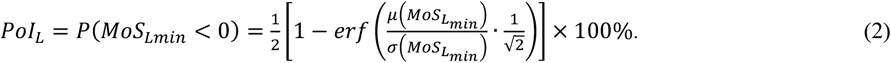

*PoI*_*L*_ thus explicitly predicts the probability a person will experience an unstable step: e.g., a *PoI*_*L*_ = 10% implies they will likely take 10 unstable (*MoS*_*Lmin*_ < 0) steps for each 100 steps they take.

### Statistical Analyses

Because the straight (STR) and winding (HIF) walking paths can be treated as distinct tasks [15, 19] we analyzed them separately. For each path, we applied two-factor (Environment × Contrast) mixed-effects analyses of variance (ANOVA) with repeated measures to test for differences between conditions for each dependent measure: % Stepping Errors, *μ*(*MoS*_*Lmin*_), *σ*(*MoS*_*Lmin*_) and *PoI*_*L*_. Environment and Contrast were fixed factors, Participant was a random factor, and the two trials served as repeated measures. For *σ*(*MoS*_*Lmin*_), to satisfy normality assumptions, we first log-transformed these data. For % Stepping Errors and *PoI*_*L*_, because multiple trials yielded resulting values equal to 0, we first added a small constant (0.001) before we log-transformed these measures. When we found significant interaction effects, we performed Tukey’s pairwise post-hoc comparisons to test for specific differences between path contrasts (High, Low) for each environment (Rich, Sparse), and between environments for each path contrast. All statistical analyses were conducted using Minitab (Minitab, Inc., State College, PA). We calculated partial-Eta squared 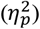 in R to evaluate effect sizes, with small effects ≥ 0.01, medium effects ≥ 0.06, and large effects ≥ 0.14 [36].

## RESULTS

### Stepping Errors

On the Straight paths (Fig. 3A), participants took more steps off the Low Contrast than High Contrast paths (p < 0.001; Table 1). On the Winding paths (Fig. 3B), participants took more steps off the Low Contrast path + Sparse environment condition than they took off either the High Contrast path + Sparse environment combination, or the Low Contrast path + Rich environment combination, (p ≤ 0.006, Table 1).

**Table 1.**
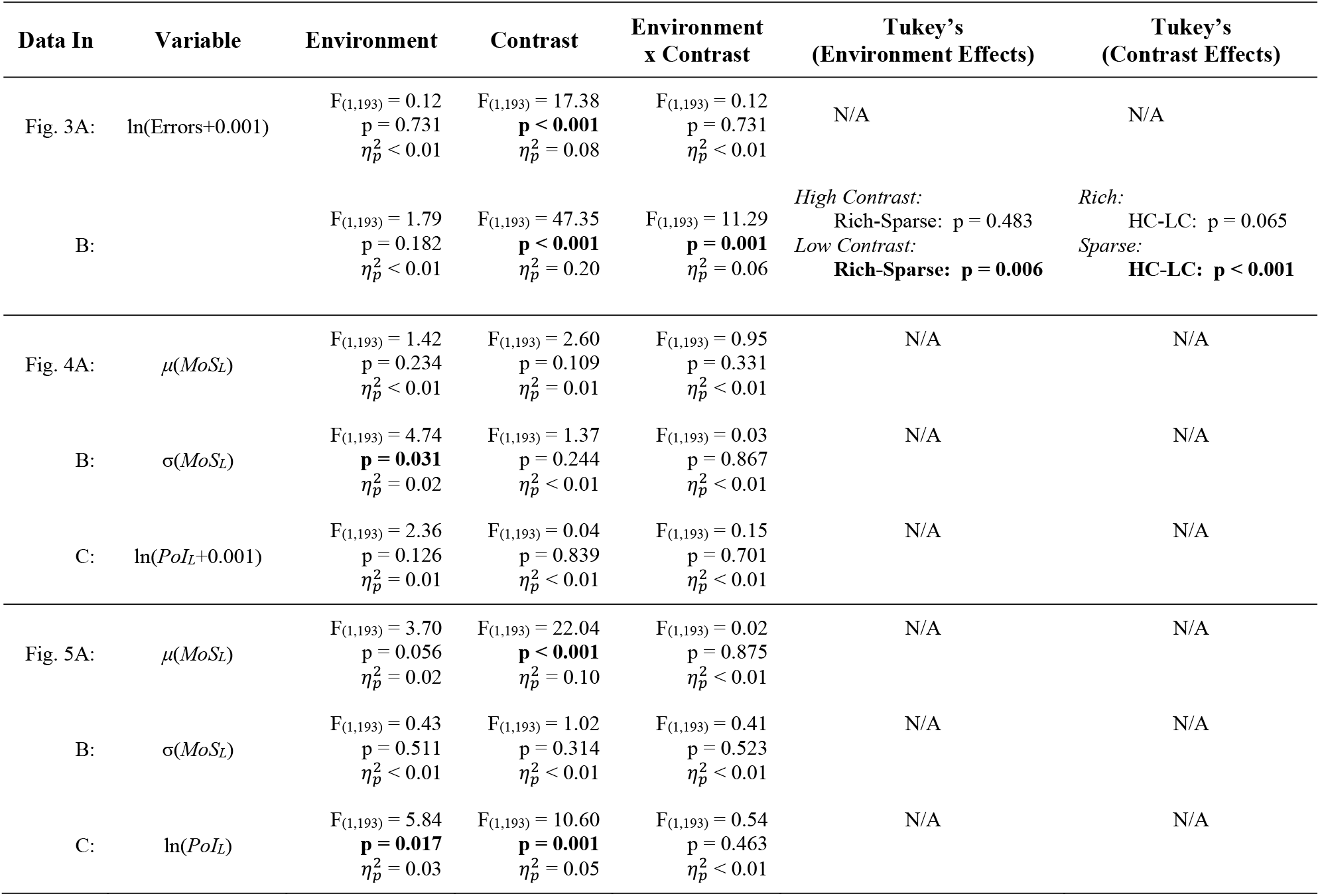
Statistical results for differences between path Environments (Rich, Sparse) and path Contrasts (High, Low) for the data shown in Figs. 3–5, including: % Stepping Errors, means (*μ*) and variability (*σ*) for lateral Margin of Stability (*MoS*_*L*_), and the lateral Probability of Instability (*PoI*_*L*_) for both Straight and Winding paths. ANOVA results (F-statistics and p-values) are provided for main effects of Environment, Contrast, and Environment×Contrast. In cases where Environment×Contrast interactions were significant, conclusions were drawn from the Tukey’s pairwise comparisons results. Significant differences are indicated in bold.

| Data In | Variable | Environment | Contrast | Environment<br>x Contrast | Tukey's<br>(Environment Effects) | Tukey's<br>(Contrast Effects) |
| --- | --- | --- | --- | --- | --- | --- |
| Fig. 3A: | $\ln(\text{Errors}+0.001)$ | $F_{(1,193)} = 0.12$<br>$p = 0.731$<br>$\eta_p^2 < 0.01$ | $F_{(1,193)} = 17.38$<br>**$p < 0.001$**<br>$\eta_p^2 = 0.08$ | $F_{(1,193)} = 0.12$<br>$p = 0.731$<br>$\eta_p^2 < 0.01$ | N/A | N/A |
| B: | | $F_{(1,193)} = 1.79$<br>$p = 0.182$<br>$\eta_p^2 < 0.01$ | $F_{(1,193)} = 47.35$<br>**$p < 0.001$**<br>$\eta_p^2 = 0.20$ | $F_{(1,193)} = 11.29$<br>**$p = 0.001$**<br>$\eta_p^2 = 0.06$ | *High Contrast:*<br>Rich-Sparse: $p = 0.483$<br>*Low Contrast:*<br>**Rich-Sparse: $p = 0.006$** | *Rich:*<br>HC-LC: $p = 0.065$<br>*Sparse:*<br>**HC-LC: $p < 0.001$** |
| Fig. 4A: | $\mu(MoS_L)$ | $F_{(1,193)} = 1.42$<br>$p = 0.234$<br>$\eta_p^2 < 0.01$ | $F_{(1,193)} = 2.60$<br>$p = 0.109$<br>$\eta_p^2 = 0.01$ | $F_{(1,193)} = 0.95$<br>$p = 0.331$<br>$\eta_p^2 < 0.01$ | N/A | N/A |
| B: | $\sigma(MoS_L)$ | $F_{(1,193)} = 4.74$<br>**$p = 0.031$**<br>$\eta_p^2 = 0.02$ | $F_{(1,193)} = 1.37$<br>$p = 0.244$<br>$\eta_p^2 < 0.01$ | $F_{(1,193)} = 0.03$<br>$p = 0.867$<br>$\eta_p^2 < 0.01$ | N/A | N/A |
| C: | $\ln(PoI_L+0.001)$ | $F_{(1,193)} = 2.36$<br>$p = 0.126$<br>$\eta_p^2 = 0.01$ | $F_{(1,193)} = 0.04$<br>$p = 0.839$<br>$\eta_p^2 < 0.01$ | $F_{(1,193)} = 0.15$<br>$p = 0.701$<br>$\eta_p^2 < 0.01$ | N/A | N/A |
| Fig. 5A: | $\mu(MoS_L)$ | $F_{(1,193)} = 3.70$<br>$p = 0.056$<br>$\eta_p^2 = 0.02$ | $F_{(1,193)} = 22.04$<br>**$p < 0.001$**<br>$\eta_p^2 = 0.10$ | $F_{(1,193)} = 0.02$<br>$p = 0.875$<br>$\eta_p^2 < 0.01$ | N/A | N/A |
| B: | $\sigma(MoS_L)$ | $F_{(1,193)} = 0.43$<br>$p = 0.511$<br>$\eta_p^2 < 0.01$ | $F_{(1,193)} = 1.02$<br>$p = 0.314$<br>$\eta_p^2 < 0.01$ | $F_{(1,193)} = 0.41$<br>$p = 0.523$<br>$\eta_p^2 < 0.01$ | N/A | N/A |
| C: | $\ln(PoI_L)$ | $F_{(1,193)} = 5.84$<br>**$p = 0.017$**<br>$\eta_p^2 = 0.03$ | $F_{(1,193)} = 10.60$<br>**$p = 0.001$**<br>$\eta_p^2 = 0.05$ | $F_{(1,193)} = 0.54$<br>$p = 0.463$<br>$\eta_p^2 < 0.01$ | N/A | N/A |

**Figure 3.**
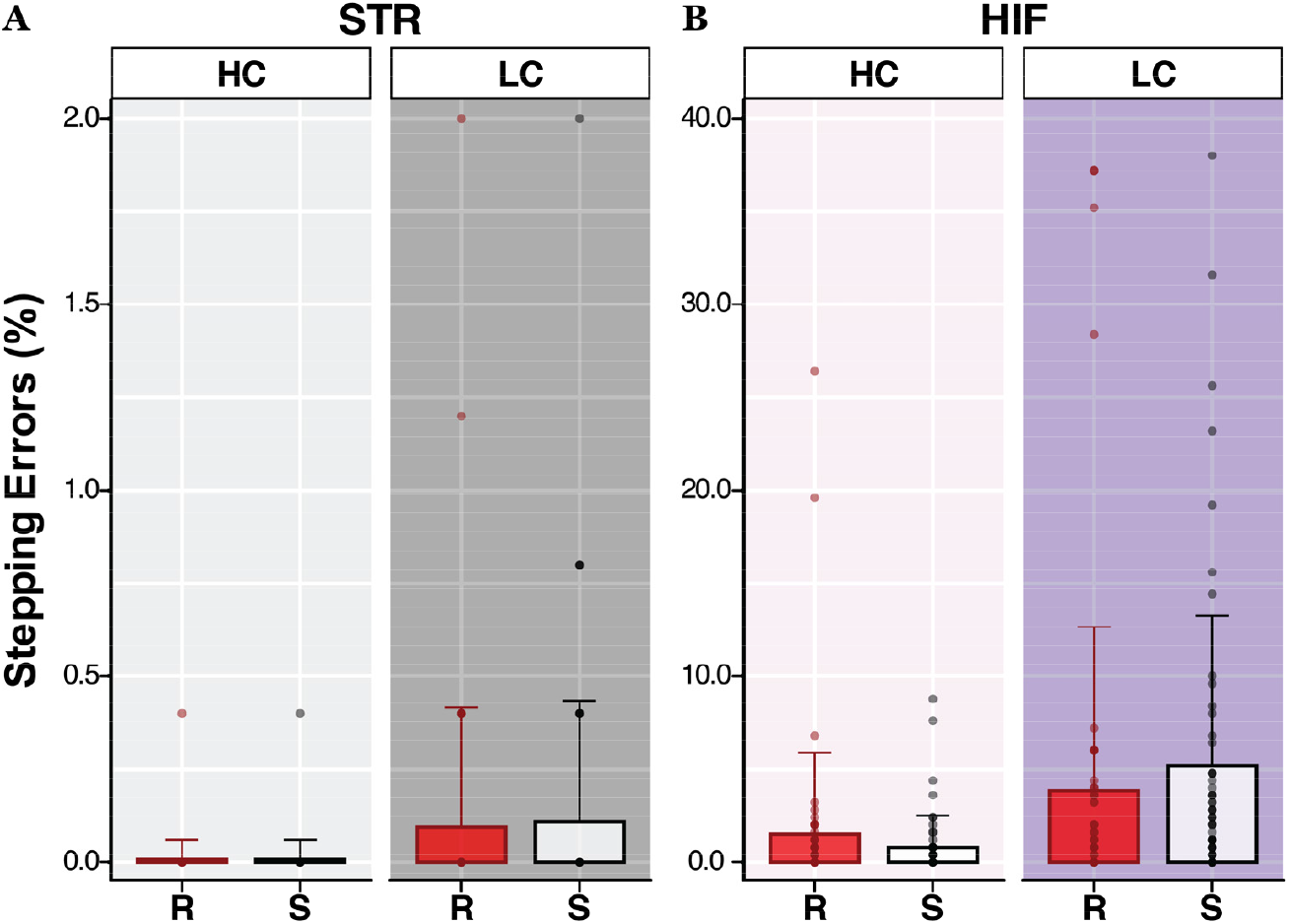
Percent stepping errors while walking on **(A)** Straight (STR), and **(B)** Winding (HIF) paths for each environment (Rich (R); filled, Sparse (S); open) and path color contrast (High (HC); light, Low (LC); dark). Bar plots represent mean values. The overlaid markers are individual data points for each participant from two separate trials. Note the vertical scale for the HIF results (**B**) [0-40%] is 20× larger than for the STR results (**A**) [0-2%]. On STR paths, participants took more steps off LC paths than HC paths (p < 0.001). On the HIF paths, this was consistent while walking in Sparse Environments, where participants also took more steps off the LC paths in comparison to the HC paths (p < 0.001). However, on LC HIF paths, participants took more steps off the path while walking in Sparse environments compared to Rich environments (p = 0.006). Results of statistical comparisons are in Table 1.

### Navigating Straight (STR) Paths

On STR paths (Fig. 4), participants exhibited margins of stability that were slightly less variable (i.e., smaller *σ*(*MoS*_*L*_); Fig. 4B) in Sparse environments (p = 0.031; Table 1). However, participants exhibited no statistically significant differences in either *μ*(*MoS*_*L*_) (p ≥ 0.109; Fig. 4A) or *PoI*_*L*_ (p ≥ 0.126; Fig. 4C) across conditions (Table 1).

**Figure 4.**
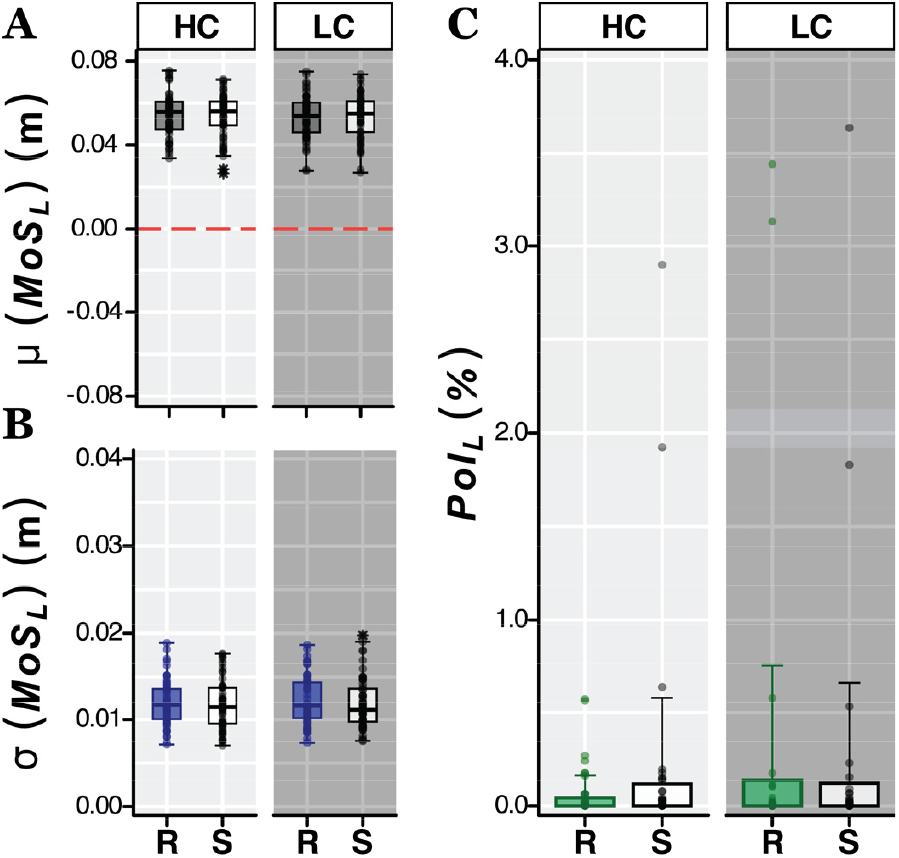
For walking on Straight (STR) paths: Lateral margin of stability **(A)** means *μ*(*MoS*_*Lmin*_), **(B)** variance *σ*(*MoS*_*Lmin*_), and **(C)** lateral probability of instability (*PoI*_*L*_) for each environment (Rich (R); filled, Sparse (S); open) and path color contrast (High (HC); light, Low (LC); dark) combination. Box plots in (A-B) show the medians, 1^st^ and 3^rd^ quartiles, and whiskers extending to 1.5 × interquartile range. Values beyond this range are shown as individual asterisks. The overlaid markers are individual data points for each participant from two separate trials. Data in (C) are plotted in the same manner as described in Fig. 3. People exhibited less variance, *σ(MoS*_*L*_), in Sparse environments (p = 0.031) but similar variance between path contrasts (p = 0.244). Results of statistical comparisons are in Table 1.

### Navigating Winding (HIF) Paths

Walking on the winding (HIF) paths was substantially more challenging than on the straight (STR) paths. On the HIF paths, participants made substantially more stepping errors (Fig. 3B vs. 3A), and exhibited margins-of-stability that were smaller on average (*μ*(*MoS*_*L*_); Fig. 5A vs. 4A), but far more variable (*σ*(*MoS*_*L*_); Fig. 5B vs. 4B), leading to much greater likelihood of instability (*PoI*_*L*_; Fig. 5C vs. 4C). These large differences replicate qualitatively similar differences found in our prior work [19].

**Figure 5.**
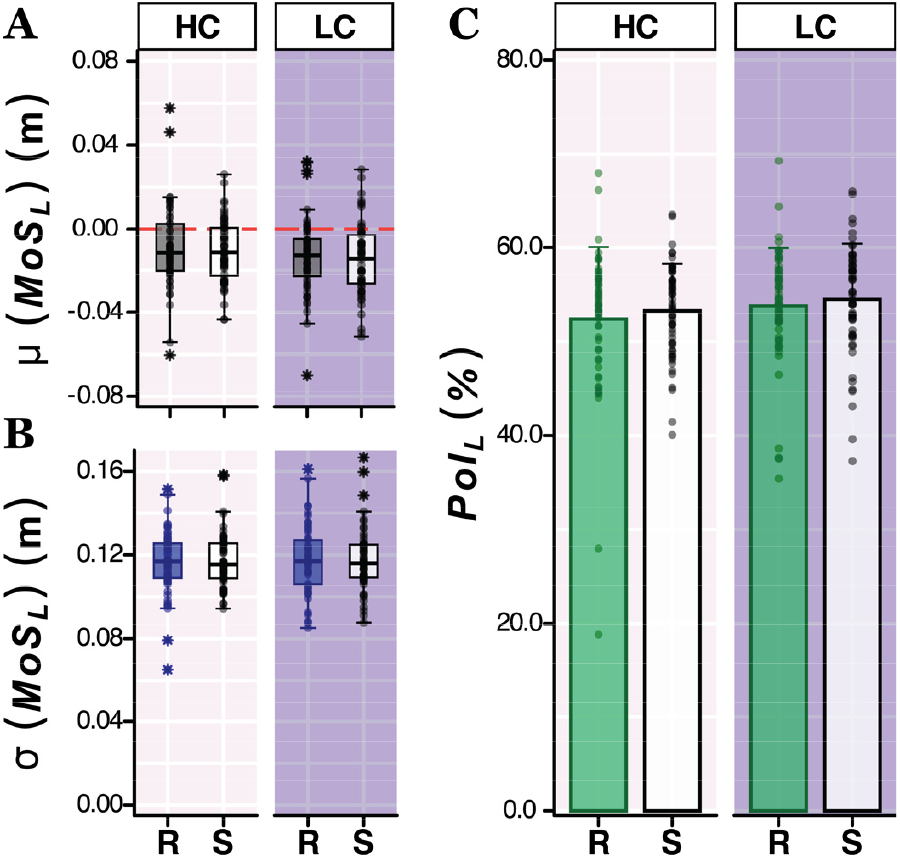
For walking on Winding (HIF) paths: Lateral margin of stability **(A)** means *μ*(*MoS*_*Lmin*_), **(B)** variance *σ*(*MoS*_*Lmin*_), and **(C)** lateral probability of instability (*PoI*_*L*_) for each environment (Rich (R); filled, Sparse (S); open) and path color contrast (High (HC); light, Low (LC); dark) combination. Data are plotted in the same manner as described in Fig. 4. Participants *μ(MoS*_*L*_) trended toward being smaller while walking in Sparse environments (p = 0.056) and were smaller on LC paths (p < 0.001). Participants exhibited higher *PoI*_*L*_ while walking through Sparse environments (p = 0.017) and while walking on Low Contrast paths (p = 0.001). Results of statistical comparisons are in Table 1.

On HIF paths (Fig. 5), participants exhibited significantly smaller *μ*(*MoS*_*L*_) (Fig. 5A) on Low Contrast paths (p < 0.001; Table 1), but only a trend in Sparse environments (p = 0.056; Table 1). Participants exhibited significantly higher *PoI*_*L*_ (Fig. 5C) both on Low Contrast paths (p = 0.001; Table 1) and in Sparse environments (p = 0.017; Table 1).

## DISCUSSION

Walking involves navigating visually complex environments [3, 4]. This requires visual-motor coordination [7, 20] to safely guide foot placement and maintain lateral stability [9, 37]. However, being adaptative also inherently heightens risk of instability [2, 8]. Here, we investigated how changes in visual perceptual salience [23] affect lateral balance and stepping performance [24] during naturalistic walking tasks. By systematically manipulating environment richness and path contrast on both straight and winding paths, we quantified the interplay between visual and mechanical challenges in maintaining lateral stability. We tested two complementary hypotheses. First, that maintaining stability is primarily governed by biomechanical task objectives [6, 25]. Alternatively, that the salience of visual information influences lateral balance, affecting stability independent of the task’s biomechanical demands, either because people use vision to enhance task-relevant information [13, 26] and reduce uncertainty [27], and/or because peripheral visual information is also used to modulate locomotor behavior [28, 29].

Differences observed between walking on the straight (STR) paths (Fig. 3A & Fig. 4) versus winding (HIF) paths (Fig. 3B & Fig. 5) demonstrate that these different paths imposed substantially different balance challenges. This replicates our prior work [19] and greatly substantiates our first hypothesis. However, the visual salience manipulations within each walking task were not inconsequential. Decreasing path visual contrast induced more stepping errors for both tasks (Fig. 3) and significantly decreased *μ*(*MoS*_*L*_) and increased *PoI*_*L*_ for the winding (HIF) walking task (Fig. 5) in particular. This supports the hypothesis that people do use visual information to reduce task-relevant visual uncertainty [13, 26, 27] about their walking path. Likewise, decreasing the visual richness of the surrounding environment induced significantly greater *σ*(*MoS*_*L*_) during straight walking (Fig. 4B) and significantly increased *PoI*_*L*_ for the winding (HIF) walking task. While these effects were fewer and somewhat smaller (i.e., smaller 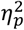) than the path contrast manipulations, they nevertheless confirm that peripheral visual information is used [28, 29] to help maintain lateral balance while walking.

On the winding (HIF) paths (Fig. 5), decreasing path visual contrast led participants to adjust their stepping. Primarily, they took slightly narrower steps that yielded smaller *μ*(*MoS*_*L*_) (p < 0.001; Fig. 5A) with no changes to variability, *σ*(*MoS*_*L*_) (p = 0.314; Fig. 5B). They likely did this to try to avoid stepping off the path (Fig. 3B), the edges of which they could not see as clearly. However, consistent with our original formulation of *PoI*_*L*_ [18] (see Eq. (2)), taking steps with smaller *μ*(*MoS*_*L*_) without correspondingly smaller *σ*(*MoS*_*L*_) directly explains their increased *PoI*_*L*_ (p = 0.001; Fig. 5C). This then suggests a possible mechanism underlying how reduced visual function might increase fall risk among older adults [11].

It is tempting to consider percent of steps taken off the path (Fig. 3) to perhaps be a proxy for risk of lateral instability (*PoI*_*L*_; Figs. 4C, 5C): i.e., intuitively, one might expect laterally unstable steps to be more likely to result in stepping errors. However, this is not obligatory. Stepping errors indicate failures in accurate foot placement (i.e., failure to keep steps on the path), whereas lateral stability measures reflect a person’s ability to maintain balance. In the present experiments, if participants strongly prioritized the walking *task*, they might sacrifice their lateral balance to minimize stepping errors (e.g., take a step within the path, when taking a step outside would have been more “stable”). Conversely, if they prioritized *balance*, they might well make more stepping errors (e.g., more wider steps off the path) to limit taking unstable steps. Here, on the straight (STR) low contrast paths, participants made more stepping errors (Fig. 3A), but exhibited no increases in *PoI*_*L*_ (p ≥ 126; Fig. 4C), consistent with prioritizing balance over task performance. On the winding (HIF) paths, participants both made more stepping errors (Fig. 3B) and took more unstable steps (Fig. 5C) when visual information was reduced (low contrast and/or sparse environment). Hence, for this more challenging task, reduced visual information salience compromised both task performance and balance control.

The reduced visual information salience had more pronounced effects on winding paths compared to straight paths, emphasizing the heightened mechanical and sensory demands of winding-path navigation. On straight paths, participants made some adjustments to their stepping (e.g., as seen in reduced *σ*(*MoS*_*L*_) in sparse environments), but these did not increase their *PoI*_*L*_ (p ≥ 0.126; Fig. 4C). In contrast, winding paths posed a greater challenge, as participants faced a higher risk of lateral instability when the quality of visual information decreased, whether due to reduced environmental richness or lower path color contrast (p ≤ 0.017; Fig. 5C).

For these young, healthy participants, many of the differences observed between the different visual manipulations were small in magnitude, even when statistically significant. Introducing sparse environments led to changes – higher *σ*(*MoS*_*L*_) on straight paths (Fig. 4B) and increased *PoI*_*L*_ on winding paths (Fig. 5C) – that were each associated with “small” [36] effect sizes (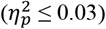). In contrast, introducing low contrast paths led to changes that ranged from “small” (*PoI*_*L*_; Fig. 5C) to “medium” (stepping errors; Fig. 3A, and *μ*(*MoS*_*L*_); Fig. 5A) to “large” (stepping errors; Fig. 3B) effect sizes [36] (Table 1). While path navigation and balance were only marginally impacted by degraded visual information, reducing path contrast elicited greater effects than reducing peripheral environmental richness.

One potential explanation for these healthy participants’ resilience could be sensory reweighting [38]. When sensory information changes, the central nervous system can prioritize more reliable sources of sensory input and adapt postural strategies accordingly [21]. Thus, our results may indicate that young people can quickly and efficiently reweight sensory inputs to maintain lateral balance, even in highly altered visual contexts. Individuals with reduced visual acuity or contrast sensitivity, such as older adults [21], could perhaps be more strongly affected by these visual manipulations.

This study extends our understanding of the sensorimotor integration processes that underpin stability during walking. Even healthy adults exhibit reduced stepping accuracy when navigating even straight low-contrast paths, with stepping errors further exacerbated on winding paths in sparse environments. These increases in stepping errors on low-contrast paths demonstrate why path boundary visibility is important for accurate foot placement [7]. Overall, our findings underscore the importance of visual salience in ensuring safe navigation and suggest that interventions focusing on sensory reweighting strategies or visuomotor training may help enhance mobility in populations at greater risk of instability.

## CRediT Author Statement

**Anna C. Render**:Conceptualization, Data Curation, Formal Analysis, Investigation, Methodology, Software, Validation, Visualization, Writing – Original Draft, Writing – Review & Editing.

**Tarkeshwar Singh**:Conceptualization, Formal Analysis, Methodology, Supervision, Writing – Review & Editing.

**Joseph P. Cusumano**:Conceptualization, Formal Analysis, Funding Acquisition, Methodology, Project Supervision, Writing – Review & Editing.

**Jonathan B. Dingwell**:Conceptualization, Data Curation, Formal Analysis, Funding Acquisition, Methodology, Project Administration, Resources, Supervision, Validation, Writing – Review & Editing.

## DATA AVAILABILITY

Data from this study are posted on DataDryad: https://doi.org/10.5061/dryad.7sqv9s56j

## ACKNOWLEDGEMENTS

The authors thank Carlie J. Hernjak and Sarah E. Overby for their contributions and technical support throughout data collections. This work was supported by NIH grants 1-R01-AG049735 and 1-R21-AG053470 (to JBD & JPC) and Sloan Foundation Grant # G-2020-14067 (to ACR).

## REFERENCES

[1] J. Acasio, M.M. Wu, N.P. Fey, K.E. Gordon, Stability-maneuverability trade-offs during lateral steps, Gait Posture 52 (2017) 171–177. 10.1016/j.gaitpost.2016.11.034.

[2] D.M. Desmet, J.P. Cusumano, J.B. Dingwell, Adaptive Multi-Objective Control Explains How Humans Make Lateral Maneuvers While Walking, PLoS Comput. Biol. 18(11) (2022) e1010035. 10.1371/journal.pcbi.1010035.

[3] M. Moussaïd, D. Helbing, G. Theraulaz, How simple rules determine pedestrian behavior and crowd disasters, Proc. Natl. Acad. Sci. USA 108(17) (2011) 6884–6888. 10.1073/pnas.1016507108.

[4] M. Nikmanesh, M.E. Cinelli, D.S. Marigold, Identifying factors that contribute to collision avoidance behaviours while walking in a natural environment, Scientific Reports 15(1) (2025) 3530. 10.1038/s41598-025-88149-3.

[5] A.E. Patla, S.S. Tomescu, M.G.A. Ishac, What visual information is used for navigation around obstacles in a cluttered environment?, Canadian Journal of Physiology and Pharmacology 82(8-9) (2004) 682–692. 10.1139/y04-058.

[6] N.S. Patil, J.B. Dingwell, J.P. Cusumano, A Model of Task-Level Human Stepping Regulation Yields Semistable Walking, J. R. Soc. Interface 21(219) (2024) 20240151. 10.1098/rsif.2024.0151.

[7] A.E. Patla, Understanding the Roles of Vision in the Control of Human Locomotion, Gait Posture 5(1) (1997) 54–69. 10.1016/S0966-6362(96)01109-5.

[8] C.E. Bauby, A.D. Kuo, Active Control of Lateral Balance in Human Walking, J. Biomech. 33(11) (2000) 1433–1440. 10.1016/s0021-9290(00)00101-9.

[9] S.M. Bruijn, J.H. van Dieën, Control of human gait stability through foot placement, J. R. Soc. Interface 15(143) (2018) 1–11. 10.1098/rsif.2017.0816.

[10] S.N. Robinovitch, F. Feldman, Y. Yang, R. Schonnop, P.M. Leung, T. Sarraf, et al., Video capture of the circumstances of falls in elderly people residing in long-term care: an observational study, Lancet 381(9860) (2013) 47–54. 10.1016/S0140-6736(12)61263-X.

[11] L.N. Saftari, O.-S. Kwon, Ageing vision and falls: a review, Journal of Physiological Anthropology 37(1) (2018) 11. 10.1186/s40101-018-0170-1.

[12] L.A. Vallis, B.J. McFadyen, Locomotor adjustments for circumvention of an obstacle in the travel path, Exp. Brain Res. 152(3) (2003) 409–414. 10.1007/s00221-003-1558-6.

[13] J.S. Matthis, J.L. Yates, M.M. Hayhoe, Gaze and the Control of Foot Placement When Walking in Natural Terrain, Curr. Biol. 28(8) (2018) 1224–1233. 10.1016/j.cub.2018.03.008.

[14] M.E. Kazanski, J.P. Cusumano, J.B. Dingwell, How Older Adults Maintain Lateral Balance While Walking on Narrowing Paths, Gait Posture 113 (2024) 32–39. 10.1016/j.gaitpost.2024.05.028.

[15] A.C. Render, J.P. Cusumano, J.B. Dingwell, Adapting Lateral Stepping Control to Walk on Winding Paths, J. Biomech. 180 (2025) 112495. 10.1016/j.jbiomech.2025.112495.

[16] A.C. Render, M.E. Kazanski, J.P. Cusumano, J.B. Dingwell, Walking humans trade off different task goals to regulate lateral stepping, J. Biomech. 119 (2021) 110314. 10.1016/j.jbiomech.2021.110314.

[17] M. Arvin, M.J.M. Hoozemans, M. Pijnappels, J. Duysens, S.M. Verschueren, J.H. van Dieën, Where to Step? Contributions of Stance Leg Muscle Spindle Afference to Planning of Mediolateral Foot Placement for Balance Control in Young and Old Adults, Frontiers in Physiology 9 (2018) 1134. 10.3389/fphys.2018.01134.

[18] M.E. Kazanski, J.P. Cusumano, J.B. Dingwell, Rethinking Margin of Stability: Incorporating Step-to-Step Regulation to Resolve the Paradox, J. Biomech. 144 (2022) 111334. 10.1016/j.jbiomech.2022.111334.

[19] A.C. Render, J.P. Cusumano, J.B. Dingwell, Probability of Lateral Instability While Walking on Winding Paths, J. Biomech. 176 (2024) 112361. 10.1016/j.jbiomech.2024.112361.

[20] W.H. Warren, B.A. Kay, W.D. Zosh, A.P. Duchon, S. Sahuc, Optic Flow is Used to Control Human Walking, Nat. Neurosci. 4(2) (2001) 213–216. 10.1038/84054.

[21] F.B. Horak, C.L. Shupert, A. Mirka, Components of postural dyscontrol in the elderly: A review, Neurobiology of Aging 10(6) (1989) 727–738. 10.1016/0197-4580(89)90010-9.

[22] D.S. Marigold, A.E. Patla, Visual information from the lower visual field is important for walking across multi-surface terrain, Exp. Brain Res. 188(1) (2008) 23–31. 10.1007/s00221-008-1335-7.

[23] D. Caduff, S. Timpf, On the assessment of landmark salience for human navigation, Cognitive Processing 9(4) (2008) 249–267. 10.1007/s10339-007-0199-2.

[24] A.A. Black, J.A. Kimlin, J.M. Wood, Stepping accuracy and visuomotor control among older adults: effect of target contrast and refractive blur, Ophthalmic and Physiological Optics 34(4) (2014) 470–478. 10.1111/opo.12141.

[25] J.B. Dingwell, J.P. Cusumano, Humans Use Multi-Objective Control to Regulate Lateral Foot Placement When Walking, PLoS Comput. Biol. 15(3) (2019) e1006850. 10.1371/journal.pcbi.1006850.

[26] M.H. Tong, O. Zohar, M.M. Hayhoe, Control of Gaze While Walking: Task Structure, Reward, and Uncertainty, Journal of Vision 17(1) (2017) 28–28. 10.1167/17.1.28.

[27] F.J. Domínguez-Zamora, D.S. Marigold, Motives driving gaze and walking decisions, Curr. Biol. 31(8) (2021) 1632-1642.e4. 10.1016/j.cub.2021.01.069.

[28] C. Vater, B. Wolfe, R. Rosenholtz, Peripheral vision in real-world tasks: A systematic review, Psychon. Bull. Rev. 29(5) (2022) 1531–1557. 10.3758/s13423-022-02117-w.

[29] D.S. Marigold, Role of Peripheral Visual Cues in Online Visual Guidance of Locomotion, Exercise and Sport Sciences Reviews 36(3) (2008) 145–151. 10.1097/jes.0b013e31817bff72.

[30] J.A. Zeni, J.G. Richards, J.S. Higginson, Two simple methods for determining gait events during treadmill and overground walking using kinematic data, Gait Posture 27(4) (2008) 710–714. 10.1016/j.gaitpost.2007.07.007.

[31] J.B. Dingwell, A.C. Render, D.M. Desmet, J.P. Cusumano, Generalizing Stepping Concepts to Non-Straight Walking, J. Biomech. 161 (2023) 111840. 10.1016/j.jbiomech.2023.111840.

[32] A.L. Hof, The ‘extrapolated center of mass’ concept suggests a simple control of balance in walking, Hum. Mov. Sci. 27(1) (2008) 112–125. 10.1016/j.humov.2007.08.003.

[33] A.L. Hof, M.G.J. Gazendam, W.E. Sinke, The condition for dynamic stability, J. Biomech. 38(1) (2005) 1–8. 10.1016/j.jbiomech.2004.03.025.

[34] K.L. Havens, T. Mukherjee, J.M. Finley, Analysis of biases in dynamic margins of stability introduced by the use of simplified center of mass estimates during walking and turning, Gait Posture 59 (2018) 162–167. 10.1016/j.gaitpost.2017.10.002.

[35] C. Curtze, T.J.W. Buurke, C. McCrum, Notes on the Margin of Stability, J. Biomech. 166 (2024) 112045. 10.1016/j.jbiomech.2024.112045.

[36] J. Cohen, Statistical power analysis for the behavioral sciences, 2nd ed., L. Erlbaum Associates, Hillsdale, N.J., 1988. 10.4324/9780203771587.

[37] C.D. MacKinnon, D.A. Winter, Control of Whole Body Balance In The Frontal Plane During Human Walking, J. Biomech. 26(6) (1993) 633–644. 10.1016/0021-9290(93)90027-C.

[38] L. Assländer, R.J. Peterka, Sensory reweighting dynamics in human postural control, J. Neurophysiol. 111(9) (2014) 1852–1864. 10.1152/jn.00669.2013.

